# Possible origins of kombucha in spontaneous fermentation

**DOI:** 10.1101/2025.07.17.665272

**Authors:** Caroline Isabel Kothe, Kim Wejendorp, Jacob Agerbo Rasmussen, Sarah Mak, Joshua D. Evans

## Abstract

Although the microbial composition and functional attributes of kombucha have been extensively studied, the origins and mechanisms of the formation of their biofilm (Symbiotic Culture of Bacteria and Yeasts, or SCOBY) remain speculative. Based on historical reports and the concept of community coalescence, this study establishes a proof-of-concept, demonstrating how a kombucha-like biofilm can form spontaneously from microbes associated with plants, insects, and humans in a sweetened tea medium. Metagenomic analyses revealed microbial dynamics during fermentation, uncovering shifts in community composition driven by acetic acid bacteria, lactic acid bacteria, and yeasts. Comparative analyses with existing fermented beverages demonstrated microbial similarities to kombucha metagenomes. Pangenomic analyses focused on *Fructobacillus evanidus*, a species present in both the bee microbiome and our spontaneous fermentation. The MAGs of *F. evanidus* from the fermented beverage showed evidence of genetic streamlining, characterized by the loss of genes essential for survival in shifting and highly competitive environments, such as the bee gut, and adaptation to the more stable conditions of fermentation. This study offers insights into the origins of kombucha, integrating historical, microbiological, ecological and evolutionary perspectives to answer the question ‘Is this a kombucha?’, and gesturing toward further physical, chemical and sensory dimensions that could be explored.

## Introduction

There is currently a significant increase in consumer demand for healthier and more sustainable food options. As part of this trend, the popularity of fermented plant-based foods and beverages is expected to continue to grow. One such product is kombucha, a sour and slightly sweet beverage produced through the fermentation of a sweetened infusion of *Camellia sinensis* tea leaves by a Symbiotic Culture of Bacteria and Yeast (SCOBY). According to market research ^1^, the kombucha market is projected to expand at an annual growth rate of 15.6% from 2022 to 2030. This beverage is often perceived as a healthier alternative to traditional sugary beverages, and has become more popular due to its reported probiotic properties, which are associated with improved gut health and boosting the immune system ^2^.

In the past decade, many scientific studies of kombucha microbial community composition have emerged, using mainly culture-independent methods and showing a symbiosis of acetic acid bacteria (AAB), yeasts, and sometimes lactic acid bacteria (LAB) ^3–5^. Some AAB species present in kombucha are known to produce cellulose, forming the biofilm—also known as a ‘mother’ or pellicle—that is used as a starter culture for successive batches ^6^. Co-cultures of bacteria and yeasts isolated from this beverage have been developed to understand their interactions and roles in biofilm formation ^7,8^. Recent studies using minimal communities have highlighted how microbial interactions shape kombucha fermentation and metabolic functions. The co-culturing of specific yeast and AAB, such as *Komagataeibacter* spp. and *Brettanomyces* spp., has been shown to generate robust biofilms ^8^. Although AAB can independently form biofilms, the presence of yeast appears to enhance their structure and resilience ^9^. A potential explanation is that yeast break down sucrose into monosaccharides and produce ethanol, which are crucial for AAB biofilm formation ^10,11^. Previous research on kombucha has also addressed aspects such as safety, product standardization, alcohol content, metabolite profiles, and sensory attributes ^6,9,12–18^.

While both kombucha producers and scientists agree that fermenting a kombucha usually requires backslopping from an existing one, this begs the question of where kombucha came from in the first place. Despite research into its microbial ecology, there is not yet any report or hypothesis in the scientific literature of how the starter culture may have originated. The increasing popularity of kombucha, and the prevalent orthodoxy that it can only be made from an existing one, have inspired our interest in testing mechanisms for its possible origins. Our hypothesis offers a viable answer to this question, which in turn also suggests there need not have been one single originating event, but rather that kombucha could have emerged multiple times in multiple places, even based on different combinations of ingredients and different microbial species, all through similar general mechanisms.

If we turn to the historical record, the origins of kombucha are surrounded by legends. It is believed that the beverage first appeared in China during the Tsin dynasty (220 BCE), and was revered for its medicinal properties ^2^. Another story suggests that kombucha originated in Japan in 414 AD, when a Korean physician cured the digestive problems of Japanese Emperor Ingyō using fermented tea ^19^. The term ’kombucha’ may have originated from the physician’s name, Kombu, with the addition of the suffix ’cha’, the Japanese word for tea. In another legend, a Russian emperor was healed by a monk who added an ant to his tea, advising him to wait for the formation of a ’jellyfish’-like structure, which was said to have curative properties ^20^. The earliest known scientific evidence about kombucha, previously called *Medusomyces* or ’tea fungus’, was reported in 1913 by a German mycologist ^21,22^.

While the exact origin of kombucha remains uncertain, these historical records suggest that East Asia is a likely geographical source, and that kombucha possibly originated through the spontaneous fermentation of sweetened tea exposed to yeasts and bacteria under favorable climatic conditions. Using the concept of community coalescence ^23,24^, we propose that the SCOBY biofilm could have arisen spontaneously through interactions among ingredients’ microbial communities, including those from plants, human skin, and/or insects, just as the mythic ant in the tea gave rise to a ’jellyfish’-like structure.

In this study, we demonstrate that a SCOBY-like pellicle can be generated through the spontaneous fermentation of typical kombucha ingredients, like green tea and honey, by microbial communities from plant, insect, and human sources gathered from the environment. In this case, one of us (KW) selected likely sources of the desired microbes from the garden of the restaurant where he worked, seeking to demonstrate how kombucha might be assembled from common sources in the environment: honeybees as they are known carriers of yeasts ^25^, sunflowers due to their association with acetic acid bacteria ^26,27^, and human skin swabs from fermentation handlers, who frequently engage in fermentative processes and may contribute relevant microbial species. Using metagenomic approaches, we analyse how these communities developed during the fermentation process to understand the formation and microbial ecology of the biofilm. Using pangenomic approaches, we then explore potential evolutionary mechanisms for how species originating from the inocula adapted to the fermented beverage environment. Finally, we compare our data with published metagenomes of other fermented beverages to assess whether and to what degree we can understand this novel beverage as a kombucha, and suggest further physical, chemical, and sensory analyses that could offer additional perspectives on this question. Overall we believe this study provides novel insights into the possible origins and mechanisms behind the formation of the kombucha biofilm, and makes a question that has until now remained speculative more tractable for scientific investigation.

## Results

### Birthing a biofilm

To investigate our spontaneous fermentation hypothesis, we carried out two exploratory experiments in a culinary environment, in 2017 and 2018. In the first year, a range of locally available ingredients were mixed into green tea sweetened with honey to inoculate the liquid with their microbial communities, including a sunflower (*Helianthus annuus*), honeybees (*Apis mellifera*), and human skin microbes grown on plates (Assembly 1). Later, in 2018, we replicated the same experiment, and also introduced a new assemblage of ingredients (Assembly 2), including rye bread and wild beach roses (*Rosa rugosa*; **Fig. 1A**). Both iterations produced similar biofilms to those of kombucha (**Fig. 1B & Fig. S1A**), and beverages with similar tastes (informal tasting, *no data shown*). Assembly 2 was not initially designed to form a biofilm, but rather to make kvass, a bread-based fermented beverage from Northeastern Europe, with the added flavor of beach roses. The addition of the beach rose petals during fermentation unexpectedly led to the formation of a kombucha-like biofilm. This serendipitous event shows how unintended microbial interactions in fermentation can lead to the emergence of spontaneous biofilm formation, and may thus also have done so in the past. These results motivated us to characterise the microbial community composition of these spontaneously formed biofilms and beverages (metadata in **Table S1**), and to see how they compared to existing kombuchas.

**Fig. 1.**
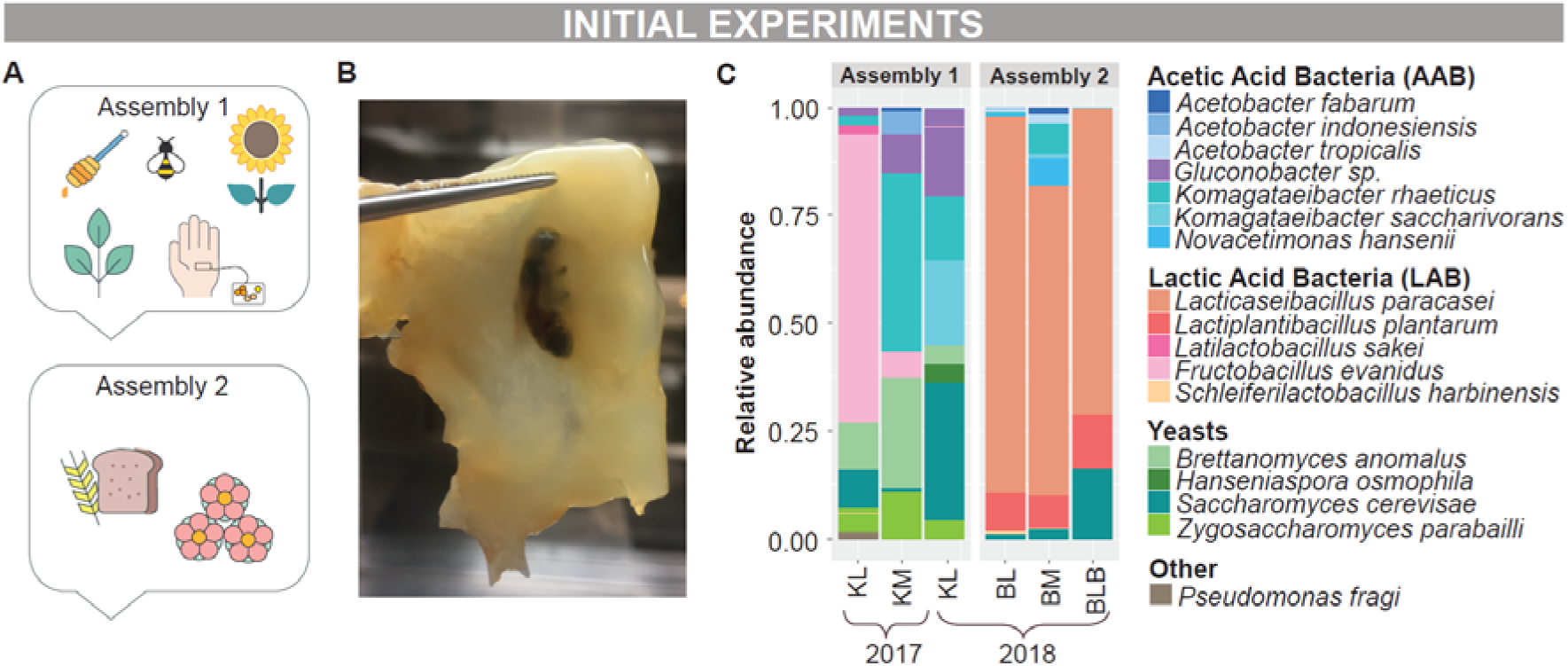
Preliminary results of spontaneous fermented beverages produced in 2017 and 2018. A. Assemblies used in the fermentation process. B. Biofilm formation from Assembly 1 in 2017, where a honeybee is visible. C. Species-level taxonomy of preliminary samples, assigned using the ychF marker gene. Relative abundance of species is calculated based on sequence coverage of assembled reads. ‘K’ indicates the beverage formed from Assembly 1 (referred to as ‘Kimbucha’), and ‘B’ that from Assembly 2 (referred to as ‘Breadbucha’). ‘L’ represents the liquid beverage, and ‘M’ the biofilm (commonly referred to as a ‘Mother’). ‘BLB’ is the liquid of the ‘Breadbucha’, bottled and stored at 5°C for one week.

All our preliminary samples presented similar microorganisms at the family level to those found in existing kombuchas: Acetobacteraceae (AAB), Lactobacillaceae (LAB), and Pichiaceae and Saccharomycetaceae (yeasts) (**Table S2**). However, the proportion of these taxonomic groups varied according to the assembly method. For Assembly 1, there was a predominance of AAB and yeast, while for Assembly 2, a predominance of LAB species (**Fig. 1C**).

These repeatable preliminary observations motivated us to conduct a more extensive and systematic experiment in a scientific environment to identify the factors shaping the microbial ecology and biofilm formation. In this experiment we sought to ask: (i) How does the microbial community develop over time? (ii) Do the raw ingredients contribute to this community and the biofilm formation, and if so, how? (iii) How microbiologically similar are our spontaneous beverages to existing kombuchas and other fermented beverages?

To address these questions, we developed an experimental design based on Assembly 1, using sunflowers, honeybees, and human hand microbes to inoculate green tea sweetened with honey. This new design included batch replicates, sampling the microbial community’s progression over time, and separate microbial analyses of each ingredient and inoculum.

### Changes in microbial community composition during fermentation

We conducted this experiment by assembling sunflowers, honeybees, and skin microbial cultures in honey-sweetened green tea (**Fig. S1 B&C**), in three separate batches (**Fig. 2A**). Initially, the pH of the mixture was 5.8, and the Brix value was 12. The fermentation process took place at room temperature (20±2°C). After one week, we observed a decline in pH and Brix to 3.3±0.19 and 9.6±0.1 (**Table S1**, sheet ‘pH’, sample ‘S0a’), respectively, across all three batches, indicating fermentation.

**Fig. 2.**
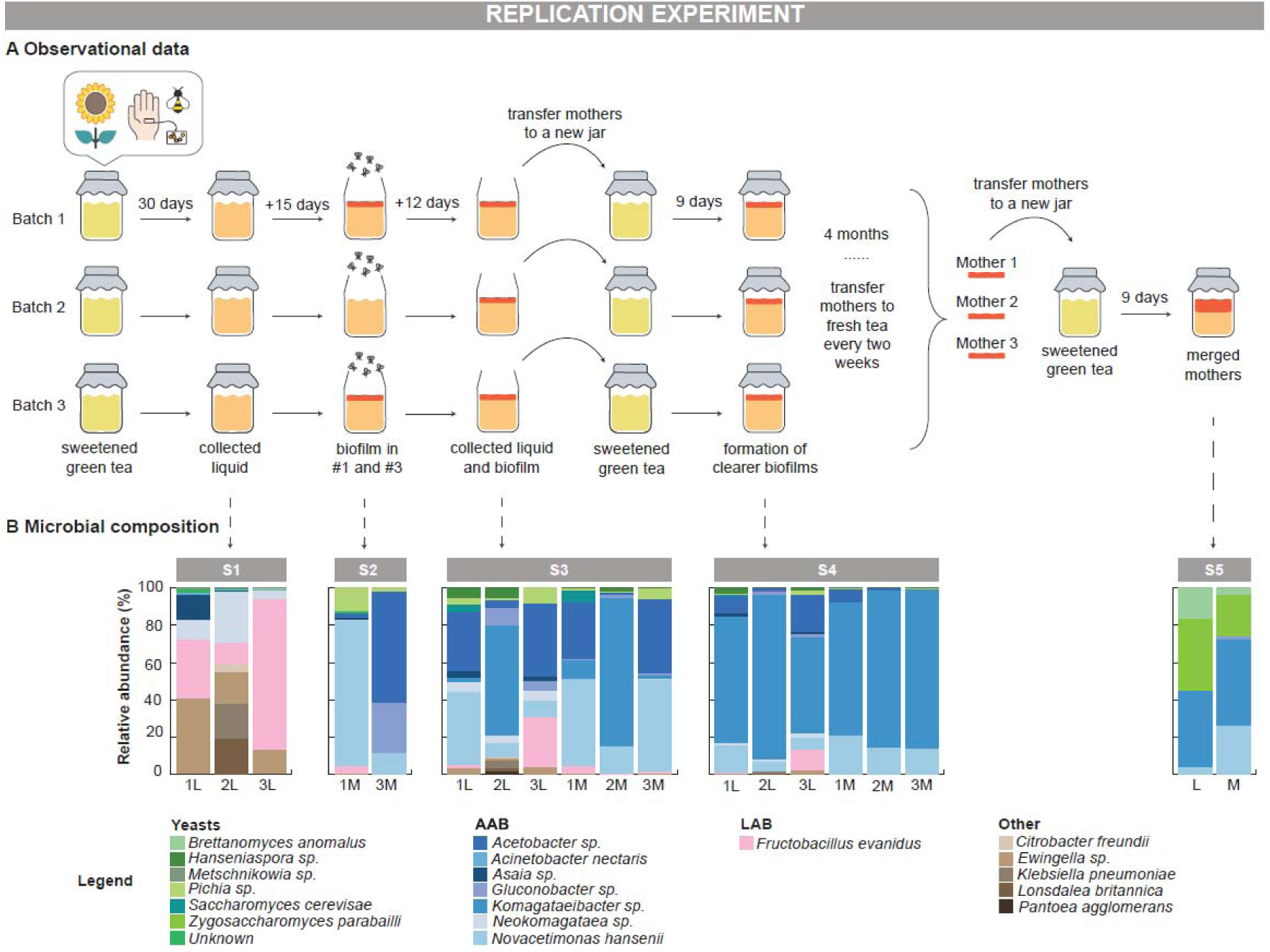
Ecological succession of microbial communities in spontaneous kombucha fermentation. A. Experimental design and timeline, including sampling points and biofilm transfers. B. The corresponding microbial composition of each sample collected. ‘S’ and the following number indicate the Sampling point; numbers preceding ‘L’ (Liquid beverage) and ‘M’ (biofilm, commonly referred to as a ‘Mother’) represent biological replicates.

After 30 days, the spontaneous fermented beverages had developed an acetic aroma, and we collected the liquid for analysis (Sampling 1). Samples show a composition dominated by AAB and LAB, with a smaller proportion of yeast (**Fig. 2B**). The main AAB was *Neokomagataea sp.*, and the main LAB was *Fructobacillus evanidus* (**Table S2**). We also noticed a high abundance of other gram-negative bacteria not common in food and beverage fermentation, the predominant species being *Ewingella sp*. Sample S1.2L exhibited high abundances of potentially pathogenic microorganisms, such as *Klebsiella pneumoniae* (18.8%) and *Lonsdalea britannica* (19.4%; **Table S2**).

After the first sampling, the batches were left uncovered, exposing them to the surrounding environment, to mimic the original experiments. At the 45-day mark, a few fruit flies were noticed in and around all three batches, and a biofilm began forming on Batches 1 and 3. The formation of the biofilm in Batch 2 may have been impeded by several factors, including microbial competition, a lower proportion of AAB, the presence of potential pathogens, the influence of fruit flies, and/or random distribution effects among the jars. We proceeded to sample these biofilms (S2.1M and S2.3M; Sampling 2). The initial biofilms exhibited high presence of the AAB, mainly *Novacetimonas hansenii, Acetobacter* spp. and *Gluconobacter* spp., as well as two species of *Pichia* yeast, *P. kudriavzevii* and *P. membranifaciens* (**Table S2**).

After an additional 12 days (57 days in total from the start point), biofilms had formed on all three batches, and samples were collected from both the liquids and the biofilms (Sampling 3). In these samples, we again identified a high abundance of AAB, now mainly *Novacetimonas hansenii, Komagataeibacter saccharivorans* and *Acetobacter persici*, as well as the presence of *Fructobacillus evanidus* (LAB) and *Pichia kudriavzevii* (yeast) across all samples.

After this period of 57 days, the pH of the liquid was 2.99±0.08 (**Table S1**, sheet ‘pH’, sample ‘S3’). At this stage, we transferred each of the mothers to a new jar containing fresh sweetened tea. After 9 days of fermentation (66 days in total), a typical length of a kombucha fermentation when microbial metabolism and chemical changes are actively taking place, we collected samples of the liquids and the mothers (Sampling 4). In these samples, we observed a continued dominance of AAB, now mainly two species: *Komagataeibacter saccharivorans* and *Novacetimonas hansenii*.

The biofilms were then maintained and renewed every two weeks for a duration of four months, to see how stable they would remain, during which no further samples were collected. Finally, after demonstrating their stability, we combined the three mothers from each batch into one new jar containing fresh sweetened tea, to consolidate space in our laboratory and to see what would happen. After 9 days of fermentation in this new tea (186 days after the beginning of the experiment), the three mothers had grown together into one, and we collected a sample from the liquid as well as from the assembled mother (Sampling 5, **Fig. 3**). In the merged sample, we observed a dominance of *Komagataeibacter saccharivorans* and *Novacetimonas hansenii*, as in the previous sampling (Sampling 4). *K. rhaeticus* was also identified in both the liquid and the mother. Notably, other yeasts not detected in previous samplings were identified in the final kombucha, and particularly in the liquid in comparable abundance to the AAB, including *Brettanomyces anomalus* and *Zygosaccharomyces parabailii,* similar to Assembly 1 in our preliminary samples (**Fig. 1**).

**Fig. 3.**
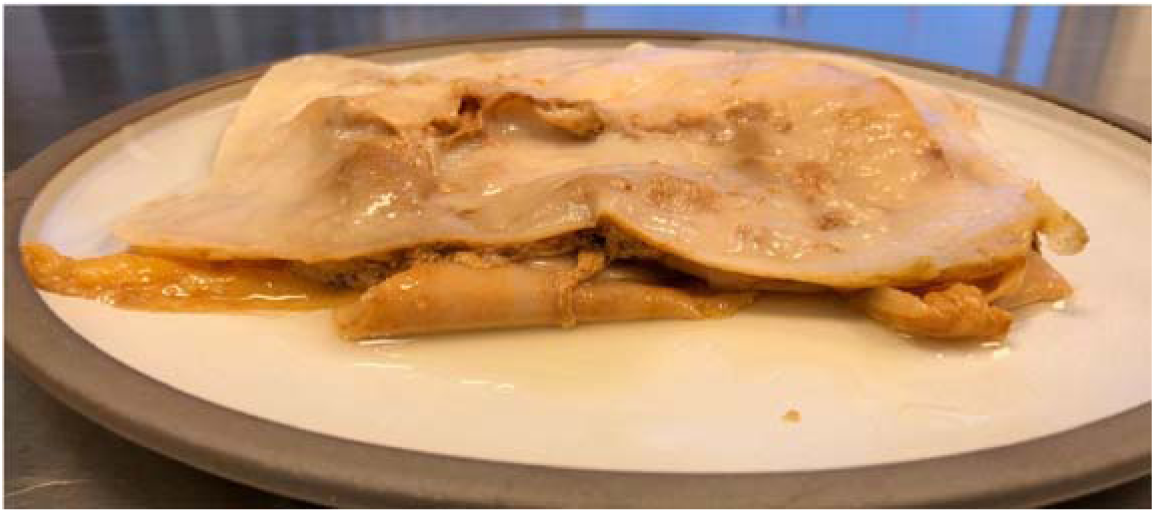
Merged mother after 186 days of experiment.

The presence of pathogens in the early stages of the spontaneous fermentation process, such as *Klebsiella pneumoniae* and *Lonsdalea britannica*, could raise safety concerns. However, as fermentation progressed and the SCOBY biofilm formed, the microbial community shifted to a safer ecology. This observation suggests that the fermentation process and the biofilm formation may have created an environment unfavorable to pathogens, reducing their abundance to below detection thresholds or eliminating them completely. We attribute this ecological succession to abiotic factors, such as low pH and nutrient availability, that are shaped by microbial metabolism, likely allowing only the most tolerant species to persist. From a food safety perspective, the final samples showed no detectable virulence factor (**Table S3**), indicating the spontaneous beverage became safer over time, and ended in a beverage that is safe to drink.

### Impact of ingredient microbial communities on spontaneous beverage ecology

In addition to tracing the development of the microbial communities over time, we investigated whether any microorganisms present on the ingredients (tea and honey) and inocula (sunflower, honeybee, and human skin swabs) contributed to the eventual microbial ecology of our spontaneous fermented beverage. To address this question, we conducted analyses of their microbiota using both culture-dependent and culture-independent methods. Of the ingredients used to prepare the sweetened tea, the green tea leaves contained diverse microbiota, including *Methylobacterium, Rhodococcus, Pantoea*, and others (**Table S4,** sheet ‘summary’). Since the tea was brewed at high temperatures, these microbiota were likely killed or inactivated and probably did not influence the fermentation process. The honey was pasteurised, and no microorganisms were detected.

The inocula, selected for their anticipated microbial richness, might have introduced microorganisms that played a role in shaping the spontaneous fermentation. However, most of the species identified in the inocula were not detected in the fermented samples. We identified *Pantoea* spp. in the sunflower (**Table S4**), as well as in the early stages of fermentation (S1 & S3; **Table S2**), indicating that they potentially came from this inoculum. Several microorganisms from the honeybees were also present in different stages of the fermentation. In addition to *Pantoea* species also found in the sunflower, the honeybees carried *Ewingella* and *Fructobacillus* spp., as well as species of the yeast *Metschnikowia*. *Pantoea agglomerans* and *Metschnikowia* spp. were only detected in the initial samples, while *Ewingella sp.* and *Fructobacillus evanidus* persisted until Sampling 4 (after 66 days of fermentation, **Table S2**). Though no species from the inocula were detected in Sampling 5, their presence in earlier samplings suggest they may have played important roles in establishing conditions suitable for a more typical kombucha ‘climax community’ to emerge.

*Fructobacillus* was also identified in our preliminary experiment from 2017 (**Fig. 1**). These observations led us to further explore the genetic relatedness of the strains of this genus across our samples through pangenomic analysis, comparing the metagenome-assembled genomes (MAGs) generated in this study (**Table S5**) with each other and against reference genomes. The *Fructobacillus* genomes available in the NCBI public database are predominantly isolated from bees and plant-associated environments (**Table S6**, sheet ‘Metrics’). In our pangenomic analysis, we identified two clades of *Fructobacillus* (**Fig. 4A**), with the MAGs from our spontaneously fermented beverage clustering closely with *F. evanidus* genomes isolated from the gut of bumblebee *Bombus lapidarius*. Although the available reference genomes come from bumblebee gut samples and our study involved honeybees, the genomic similarity suggests a potential diet-and niche-based association rather than strict host specificity. Average Nucleotide Identity (ANI) analysis showed that these genomes of *F. evanidus* shared >99% ANI, indicating a high level of genetic similarity (**Table S6**, sheet ‘ANI’).

**Fig. 4.**
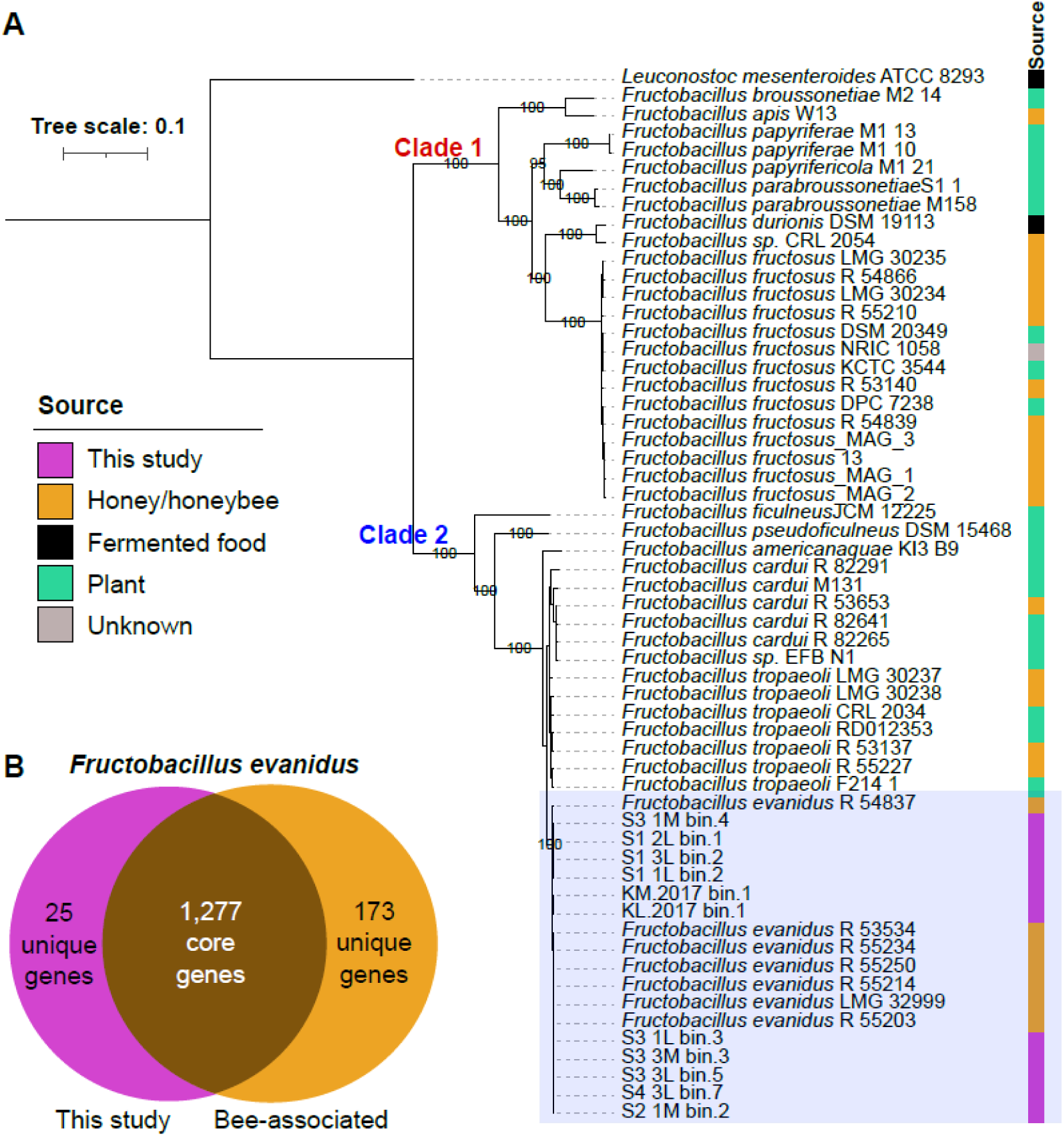
Pangenomic analysis of Fructobacillus spp. A. Phylogenetic tree of Fructobacillus species, highlighting two distinct clades. MAGs from this study, classified as Fructobacillus evanidus, are clustered within Clade 2, indicated by the blue box on the corresponding branch. B. Venn diagram showing the shared and unique gene content of Fructobacillus evanidus genomes from this study (pink) and bee-associated sources (orange).

While the average genome size of *F. evanidus* strains from bees is around 1.7 Mb, our MAGs have a notably reduced genome size of 1.6 Mb, suggesting potential gene loss in the fermentation environment. To investigate which genes differentiated the bee-associated reference genomes from our spontaneous fermentation ones, we compared their sets of core genes. Our MAGs shared 1,302 core genes among them, while the bee reference genomes shared 1,450 core genes. The two groups shared 1,277 core genes, with 173 unique to the reference genomes, and only 25 unique to our samples (**Fig. 4B**). Among the unique genes in the reference genomes, we identified 47 enzyme-related genes, which may reflect an adaptation to the bee gut environment, where these bacteria are likely to encounter more challenging and variable conditions compared to the relatively stable conditions of the fermented beverage environment (**Table S7**). In our MAGs, the unique genes included several hypothetical proteins of unknown function, as well as seven phage-related genes, which may suggest genome reduction or gene loss events.

### Comparing microbial profiles of our spontaneous beverages and existing fermented beverages

To better understand how our spontaneous beverage relates to other fermented beverages microbiologically, we performed a Principal Component Analysis (PCA) of our metagenomic data and some found in the literature (**Table S8**). Previous studies using shotgun metagenomics have provided microbial profiles of kombuchas ^3,8,17,28^, wine ^29^, vinegar ^28,30^, water kefir ^28,31,32^, lambic beer ^33^, pu-erh tea ^34^, pulque ^35^, and other fermented beverages using fruits, cereals and honey ^28^. By comparing our samples with these beverages, we assessed how much our spontaneously fermented beverage shares microbial characteristics with established profiles like kombucha or exhibits unique fermentation patterns (**Fig. 5A**).

**Fig. 5.**
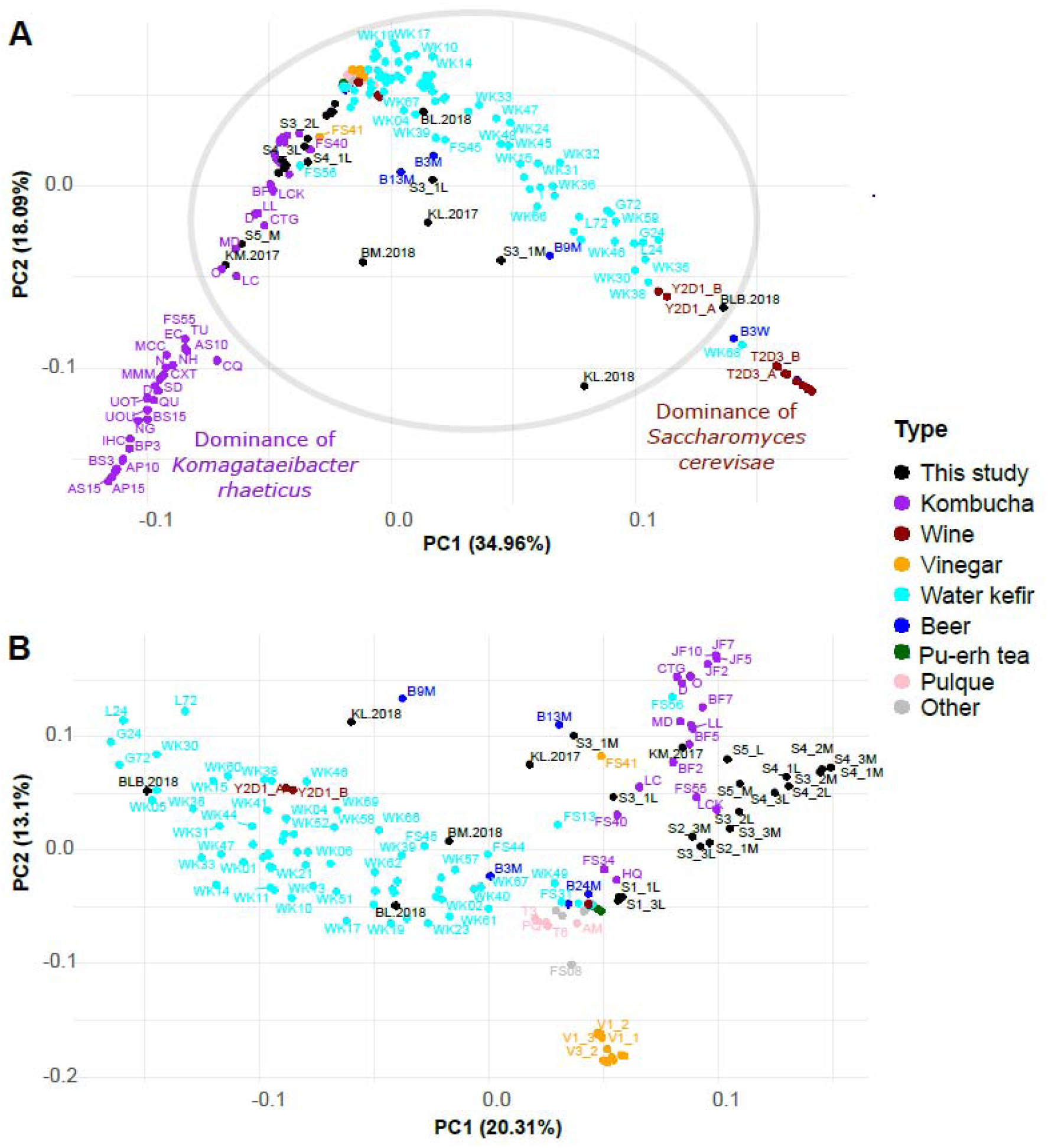
Principal Component Analysis (PCA). **A.** Plot comparing the spatial placement of our fermented beverages (black points) with other fermented beverages from the literature, based on metagenomic data. **B.** Plot showing a closer comparison of our fermented beverages with a subset of closely related beverages (excluding outliers and only considering the samples inside the circle from Fig. 5A).

The PCA shows a clear two-axis pattern, suggesting two main microbial ‘poles’ toward which these samples are orthogonally distributed. The left cluster is predominantly composed of kombucha samples, dominated by *Komagataeibacter rhaeticus*, an AAB known for cellulose production, which contributes to the physical structure of kombucha biofilm and stabilises the kombucha microbial ecosystem. The right cluster includes samples typical of wine fermentations, dominated by *Saccharomyces cerevisiae*, a yeast that converts sugars to ethanol during alcoholic fermentation. This separation reflects the fundamental differences between kombucha and wine fermentations, even when considering the third PCA component, which accounts for 10% of the variance (**Fig. S2A**). Along with the majority of our samples (black dots) are additional samples from other studies: kombucha, water kefir, vinegar, pulque, and Belgian lambic beers that have been fermented and matured for more than three months in wooden barrels, suggesting an overlap in their microbial communities. Two hypotheses may explain the nature of this overlap: (i) some of these central samples might represent more diverse microbial communities due to mixed ecologies and/or fermentation environment; and/or (ii) some might still be in an early stage of microbial development, with their communities evolving toward a more specific and stable ecology over time.

While this analysis shows the broad microbial similarities between our samples and known beverage categories, a focused PCA excluding the extreme poles revealed that our samples cluster mostly with the kombucha samples from the literature, dominated by *Brettanomyces sp.* and *Komagataeibacter sp.* (**Fig. 5B**). The ’Breadbucha’ samples cluster closer to several water kefir metagenomes, which can be explained by the presence of LAB and *Saccharomyces* species likely originating from the sourdough bread used in the preparation and/or from the surrounding environment which contained various ongoing fermentations. Most of the vinegar samples ^28,30^ clustered separately from the kombucha samples, including ours. Additional insights into microbial community differences at various fermentation stages emerged when considering the third PCA component, which accounts for 12.7% of the variance—a value comparable to PC2 (**Fig. S2B**). Our early-stage samples (Sampling 1) cluster most closely with our preliminary samples (**Fig. 1**) and other mixed fermentations, likely due to a shared presence of LAB species. The intermediate-stage samples (Samplings 2, 3 & 4) cluster in the lower-left corner of the plot (**Fig. S2B**), suggesting a transient microbial community dominated by *Komagataeibacter saccharivorans* and *Novacetimonas hansenii*. And by the end of the experiment, the final samples have again moved closer to other kombuchas.

Further supporting this progression is the presence of potentially novel species in the initial and intermediate samples of our study. The MAGs recovered from our samples suggest the presence of three potentially novel species belonging to the genera *Asaia*, *Ewingella*, and *Neokomagataea* (**Table S5**), highlighting the broad and partially unmapped microbial diversity that emerged during the early stages of spontaneous fermentation.

## Discussion

Determining whether our spontaneously fermented beverage is a kombucha is a complex question that must be approached from different perspectives. This question must take into account not only microbial profile, but also historical context, ecological and evolutionary dynamics, physical properties of the biofilm, volatile compound profiles, sensory attributes, and cultural acceptance. Each of these perspectives offers different insights into what kombucha is, highlighting the value of a holistic approach to understanding and characterising fermented foods and beverages. In this discussion, we address our central question—*Is it a kombucha?*—by presenting a microbiology-focused proof-of-concept based on the beverage’s historical context, but also exploring additional dimensions that contribute to its classification as kombucha.

### Historical context

Although there are suggestions that kombucha has been consumed for thousands of years in different parts of the world—mostly associated with health claims ^36^ and therapeutic properties ^37^—its exact origin remains uncertain. In this study, we offer insights on the possible origins of kombucha by recreating a sour, sweet, bubbly beverage, safe for consumption and complete with SCOBY-like biofilm, through spontaneous fermentation. This discovery raises questions about the origins of kombucha, and especially its biofilm. Might it have emerged in the past through a similar interaction of ingredients and fermentation conditions? Many traditional ferments likely emerged through such fortuitous accidents and were subsequently refined over generations ^38^. For example, the origin of cheese is believed to have been accidental, possibly discovered when milk was stored in containers made from animal stomachs. Rennet, an enzyme found in the stomachs of ruminant animals, would cause the milk to coagulate, separating it into curds and whey ^39^. Similarly, vinegar has its roots in the souring of wine, and was produced through a slow, natural and spontaneous fermentation process driven by AAB ^40^. It is possible, if not likely, that a similar process happened for kombucha—as illustrated by the serendipitous emergence of the Breadbucha (Assembly 2) after adding rose petals to a kvass. While it may not be possible ever to conclusively discern these origins, our replicable proof-of-concept shows one mechanism by which the spontaneous emergence of the biofilm would have been possible, even with different ingredients and under different fermentation conditions.

These different methods raise the question of how these beverages relate to kombucha *vs* jun, and whether the latter two are different microbiologically. Some fermenters distinguish jun (or xun) as a kombucha-like beverage traditionally made with green tea and honey, whereas kombucha is typically associated with black tea and sugar. Both black and green tea are made with leaves of *Camellia sinensis*, but undergo different processing methods that influence their polyphenol composition, acidity, and flavor profile ^41^. Sucrose, a disaccharide composed of glucose and fructose, requires microbes like *Brettanomyces* to produce invertase for hydrolysis before fermentation is possible ^42^, while honey naturally contains around 70% of monosaccharides, making sugar readily available for fermentation by yeasts and acetic and/or lactic acid bacteria ^43^. Despite this practical distinction among some fermenters and consumers, kombuchas fermented with black or green tea do not appear to differ significantly in their bacterial composition ^2,44,45^. Yeast composition may differ slightly depending on the tea type ^44^, although these differences are not consistently characterized in the literature. Furthermore, while jun is often claimed to have originated in Tibet or northern China, Katz argues this narrative may be a marketing fabrication, as no historical records mention it in Tibetan or Himalayan fermentations ^46^. Based on their microbiological similarity and the lack of historical records for jun, we believe it most prudent to understand kombucha and jun, and by extension our spontaneous fermented beverages, as different types of kombucha, showing how kombucha-like beverages can arise independently in different regions, shaped by local ingredients, inocula, and fermentation practices.

### Microbiology, ecology & evolution

Overall our samples, especially those of the replication experiment, seem microbiologically comparable to kombuchas. The microbial communities present in our final spontaneously fermented beverages, particularly AAB, such as *Komagataeibacter* spp., and yeasts such as *Brettanomyces* spp. and *Zygosaccharomyces* spp., closely resemble those in kombucha ^3,4,28^. Early in the fermentation (Sampling 1), we observed a community dominated by *Fructobacillus evanidus*, a LAB species, and *Ewingella sp*., likely introduced via the inocula. As fermentation progressed, these early colonizers decreased, giving way to a transient intermediate community, dominated by AAB and *Pichia kudriavzevii* as the main fungal species (**Table S2**). Over time, this community shifted to a more balanced composition, with a reduced dominance of AAB and an increased proportion of yeasts— microbiologically comparable to that of kombucha, suggesting a convergence in microbial succession (**Fig. 5**).

The PCA generated using metagenomic data suggests a clear difference between vinegar and kombucha microbial ecologies (**Fig. 5B**). While both fermentations rely on AAB, in vinegar they tend to dominate, often with minimal yeast activity remaining, whereas kombucha typically maintains a more balanced coexistence of yeasts and AAB. These ecological differences correspond to differences in production technique. Vinegar production typically has two stages of fermentation: an initial alcoholic fermentation, occurring anaerobically, where yeasts convert sugars into ethanol, followed by acetic fermentation in an aerobic environment, where AAB oxidise ethanol into acetic acid ^50^. Kombucha, while it can have an additional anaerobic fermentation in bottle to produce bubbles, only requires a one-stage aerobic fermentation, with yeast and bacteria metabolising concurrently.

Kombucha also typically ferments faster than vinegar, at least if both are made using traditional passive methods. When brewing kombucha, fermentation is typically stopped after about a week to have a beverage that is effervescent, pleasantly sour, and somewhat sweet, according to the brewer’s taste. If fermentation continues, all the sugar in the kombucha will be converted into acetic acid. In practice, this means that kombucha that ferments for too long will eventually turn into a kind of vinegar, and if this happens it is commonly used as such, more as a seasoning in cooking than as a beverage ^46^.

Beyond the question of whether this is a kombucha, there are additional insights we can gain from studying microbial evolution in this system on how environmental microbes adapt to the fermentation niche. Here we focus on *Fructobacillus evanidus*, a species detected in our fermented samples that is also known to inhabit the gut of bees. The differences in gene content between fermentation-associated MAGs and bee-associated genomes provide insights into microbial adaptation. Bee-associated genomes appear to retain additional genes that equip them to survive in a more variable and potentially harsher environment, characterised by exposure to parasites, pathogens, pests, pesticides and climatic changes ^51–53^, as well as diverse food sources, including pollens and nectars that vary seasonally over the year ^54,55^. These genes may encode enzymes that enhance nutrient acquisition, stress response, and detoxification processes, which are crucial for survival in the dynamic and challenging conditions of the bee gut ^56^. In contrast, the relatively stable fermentation environment may have allowed our MAGs to lose certain genes that are no longer essential for survival, resulting in genetic streamlining, or genome reduction. This adaptation reflects a shift toward metabolic efficiency in a more predictable habitat, where certain stress responses and metabolic functions become less critical. A classic example of this phenomenon can be observed in domesticated *Lactococcus* species. Originally found in plant environments, *Lactococcus* strains introduced into the dairy fermentation ecosystem have lost the ability to synthesise various amino acids and degrade plant-derived sugars, as a result of adapting to the conditions of milk ^57,58^.

Such fermentation microorganisms are the result of thousands of years of genotypic and phenotypic modification throughout human interaction with food, leading to what we now recognise as microbial domestication ^59^. Ancestral practices, such as backslopping, have played an important role in the long-term transmission and gradual modification of microbial populations. These populations, isolated from their initial cultures and placed in new conditions, gradually adapt to them and become genetically differentiated ^60,61^. A key aspect in domestication is this shift from microbes’ natural environment—often a hostile terrain characterised by intense competition and sporadic nutrient availability—to a more stable, nutrient-rich, human-engineered environment ^59^, with reduced competition and conditions that favor desirable microbiological traits and corresponding genome optimisation ^62^. The genetic adaptation of *Fructobacillus evanidus* we witness in our ‘kombuchas’ from the bee gut to the fermented beverage environment is one such example that, if perpetuated, could lead to a comparable domestication. And while it may seem strange that novel fermentation communities could coalesce from microbes associated with plants, insects, other animals, or even humans, such holobiotic exchanges of microbes as a generator of fermentation microbial ecologies seems much more the rule than the exception ^63^.

Overall, the kombucha-like microbiology observed in our fermented beverages provides evidence for the possible origins of kombucha in spontaneous fermentation based on community coalescence.

### Physical, chemical & sensory dimensions

Although our study focused on microbiological aspects, we recognize the importance of other dimensions—such as the physical structure of the biofilm, volatile compound profiles, and sensory attributes—in defining what makes a beverage ’kombucha-like’. We therefore discuss these dimensions and the extent of knowledge on them in the literature to further contextualise our findings and broaden the discussion about what characterises kombucha.

The physical structure and properties of the biofilm can be one parameter for defining kombucha and differentiating it from other fermented beverages. In kombucha, the biofilm forms a thick, gelatinous film at the liquid–air interface, mainly composed of cellulose produced by the AAB ^6^. Besides AAB, yeasts play a crucial role in shaping the biofilm structure, forming pseudo-mycelium that supports the cellulose network ^9^. Other fermented beverages also form biofilms or microbial matrices, but they differ in their physical properties. For example, water kefir grains—a well-known microbial consortium used as a starter culture typically for fermented beverages based on fruits—are small, translucent, irregularly shaped aggregates of exopolysaccharides, mainly dextran and levans, forming a matrix that encapsulates a mix of LAB and yeasts ^64^. Other fermented products can also develop biofilms, as in the case of vinegar ^48^. The biofilm in vinegar, often called ’mother of vinegar,’ is typically a loose gelatinous matrix similarly composed of cellulose-producing AAB that settles at the bottom of the fermenting liquid rather than floating on top in a thick layered mat like kombucha mothers ^65^. In other fermented beverages, biofilms are not desirable, such as in beer production, where they result in unwanted flavors and aromas, turbidity, and potential microbiological instability in the final product ^66,67^. The biofilm formed in our study, with its thick, layered, mat-like structure forming at the liquid–air interface, is closest to a kombucha biofilm, and not water kefir or vinegar (**Fig. 3**).

Though kombucha is brewed with sweetened tea, typical tea aromas have not been highlighted in the characteristic aromatic profile of the final product ^68^. Recently, Wang et al.^18^ proposed potential flavor quality markers for kombucha, emphasising several key volatile compounds. These include floral alcohols (2-phenylethanol), sweet aldehydes (2,5-dimethylbenzaldehyde), acetic acid for vinegar-like notes, and esters like ethyl acetate and ethyl hexanoate, which collectively bring fruity, floral, and sweet notes. However, there is still no clear definition of the volatile compounds essential for its aroma, or even if this is something that could be standardised, given the wide variety of kombucha cultures, producers, and production methods. Future studies could apply untargeted and targeted volatilomics and consumer sensory analyses across a diverse range of kombucha samples to identify conserved aromatic signatures.

Sensory attributes, including sourness, effervescence, and flavor profiles, are currently more tractable parameters for characterising kombucha scientifically. This multisensory approach aligns with Redondo et al.’s ^70^ assertion that taste perception of soft drinks (to which kombuchas are similar in taste if not production method) involves a complex combination of visual, olfactory, gustatory, and tactile sensations. There are several ways to make kombucha, and depending on the producer and/or consumer’s preference, the fermentation time and temperature can be varied to achieve different levels of sourness, sweetness, aroma profile, and overall sensory experience. Curiously, there are limited studies on descriptive sensory attributes of kombucha tea beverages. Neffe-Skocińska et al. ^15^ found that kombucha fermented for 10 days using black and green tea with 10% (w/v) sucrose at 20-30°C was described as ’tea’, ’citrus’, and ’acid’ by 16 untrained participants. Ivanišová et al. ^14^ reported a ‘pleasant, fresh, sour-fruity’ taste in kombucha fermented for seven days with black tea leaves (3% sugar) at 22°C, based on a larger panel of 60 evaluators. Some also noted a ‘vinegar taste’, which suggests a sensory overlap between kombucha and vinegar. These scientific findings substantiate common and widespread knowledge about kombucha sensory profiles known to fermenters for generations.

These findings illustrate the complex and varied sensory characteristics of kombucha beverages, which are influenced by multiple factors, including the type of ingredients, sugar concentration, fermentation temperature and duration, as well as producers’ and consumers’ sensitivities and preferences. Though we do not include a formal sensory analysis of our spontaneously fermented kombucha-like beverage in the present study, from our own tasting we can state that it exhibits the same sour-sweet, aromatic, and effervescent qualities present in other kombuchas, and when the preliminary samples were served at the restaurant where they were developed, they were accepted by guests as kombucha.

## Conclusion

To test the hypothesis that kombucha could have originally emerged through spontaneous fermentation based on community coalescence, we conducted two preliminary and one systematic replication experiment combining plant, insect, and human microbiomes in sweetened tea. All fermented successfully, yielding sour, effervescent beverages and robust SCOBYs. Based on metagenomic data, we tracked changes in microbial community composition during fermentation, assessed the impact of ingredient microbial communities on spontaneous beverage ecology, and compared microbial profiles of our spontaneous beverages with existing fermented beverages. Over the course of fermentation, the microbial communities transitioned from a more heterogeneous composition influenced by the inocula to a structure more typical of kombucha, characterized by a balanced presence of yeasts and AAB. While some inocula-derived microbes entered the initial and intermediate samples, only a couple persisted, and none were detected in the final samples. They may have played important roles in stabilising the emergent kombucha ecology, giving way to more typical kombucha species as the ecology approached the kombucha ‘climax community’. Microbiologically, the resulting beverages were similar to existing kombuchas, and most similar to kombuchas than to other fermented beverages like water kefir or liquid seasonings like vinegar. The SCOBY biofilms were also most physically similar to those of kombucha than to the others, as was the sensory profile; for these we present only observational data. In addition to these different answers to the question of whether these spontaneously fermented beverages are kombuchas, we also traced the adaptation of strains of *Fructobacillus evanidus* from the bee microbiome to the fermented beverage environment, highlighting a potential instance of incipient microbial domestication. Taken together, this knowledge not only deepens our understanding of kombucha and its possible origins specifically, but also provides the basis for future research into microbial succession and adaptation across other fermented food and beverage systems.

By bringing together not only microbiology, ecology, and evolution, but history, physics, chemistry, and sensory science, these questions of kombucha’s possible origins offer rich avenues for continued interdisciplinary investigation. Further research could explore the historical, archeological, and anthropological records, as well as current cultural practices associated with kombucha, to identify any further clues about the beverage’s evolution and relation to other fermented beverages, microbiologically and culturally. Investigating physical variations in biofilm formation and structure, microbial composition across different regions and ingredients, and chemical and sensory variability could shed light on potential core microbial communities, metabolomes, and/or sensory profiles, and how they relate to other foods and beverages. More frequent sampling, including at the start of fermentation as well as when the kombuchas over-fermented and became more like vinegar, could provide finer granularity in tracking microbial succession and ecological shifts over time. Together, these investigations could elucidate the drivers that have shaped kombucha throughout its history—and, as many kombucha makers today are experimenting with a range of non-traditional ingredients and methods ^71^, continue to shape its future.

## Materials and Methods

### Initial experiment samples

The preliminary experiment was conducted in a restaurant environment, with two different assemblies.

First, we used sweetened tea inoculated with honeybees, sunflower and human skin microbiota to recreate kombucha from scratch. The experiment was performed in two consecutive years, 2017 and 2018. Green tea (Kolonial, Denmark) was brewed using 32 g of tea in 4 liters of boiling water. After cooling to room temperature, honey (Jakobsens, Good Food Group A/S, Denmark) was added and stirred in to achieve a Brix level of 8%. The mixture was then transferred to a 5-liter plastic container (Condi ApS, Denmark). Alive honeybees and sunflowers were collected from the beehives and garden of restaurant Amass (Copenhagen, Denmark). Skin samples were collected using a cotton swab from the hands of the fermenter, plated directly on PDA (Potato Dextrose Agar) plates, and incubated at 30°C for 24 h. These ingredients were subsequently added to the tea mixture and fermented at 25-30°C, covered with a synthetic-fibre cloth. The fermentation room contained various other ongoing fermentations. Fruit flies were occasionally present. The fermentation process was carried out for approximately 2-3 weeks, until the formation of a pellicle was observed. At this stage, the ‘mother’ and a portion of the fermented liquid were transferred to a fresh batch of tea and honey mixture for continued fermentation. Once a stable pellicle formed, fresh tea was provided every two weeks. The samples were collected 6 months after the stable pellicle formed.

The second assembly was originally designed as a kvass, a traditional fermented beverage made from bread. The intent was to develop a kvass with added flavors from beach rose (*Rosa rugosa*). However, during the process, an unexpected biofilm formation was observed, leading to a fermentation process different than intended. Leftover rye bread (Lille Bakery, Denmark) was toasted in an oven (EasySteam, Electric Combi Oven 10GN 1/1, Zanussi, Italy, 170°C). A mixture was prepared in a steel gastro-tray (H.W. Larsen, Denmark) containing 1 kg of toasted bread, 250 g of pilsner malt (Viking malt, Finland) and 6 liters of boiled tap water. The mixture was incubated at 60°C for 2 h to activate α-amylase in the malt, followed by 70°C for 1 h to activate β-amylase. The liquid was then strained, producing a solution of approximately 10°Brix. This liquid was further reduced on a stove (Modular Cooking Range Line EVO700, Zanussi, Italy, setting 4) until it reached a concentration of 40°Brix. From 1 kg of leftover bread, approximately 1 kg of 40°Brix syrup was obtained. A subsequent solution was prepared by combining 1 liter of 40°Brix syrup with 1 liter of tap water and 600 g of sugar (Dansukker, Denmark), and left to ferment for 5 days, covered with a synthetic-fibre cloth. Beach rose petals (50 g) were then added to enhance flavor. After 12 days, a biofilm resembling that of a kombucha formed. The biofilm was then treated as a kombucha, transferred every two weeks to fresh sweetened tea, prepared as described above (green tea with honey, 8**°**Brix). The samples were collected 6 months after the stable biofilm formed.

### Sampling of ingredients and spontaneous kombucha replication experiment

Sunflower heads and live honeybees (*Apis mellifera*) were collected from the garden of restaurant Amass (Copenhagen, Denmark) in July 2022. A swab was taken from the left and right hand of the person responsible for producing the original kombuchas (front, back, and between the fingers) and was diluted in 1 ml of 0.9% NaCl solution. This solution was homogenised, and 100 µl were poured onto four Brain Heart Infusion (BHI) agar plates. After 24 h of incubation at 30°C, three plates were prepared for the experimental kombucha batches and one for further strain characterisation. The other ingredients used were pasteurised honey (Jakobsens, Denmark), green tea (Chinese sencha, Tante T, Denmark) and tap water.

Tea (16g) was brewed in 2 liters of boiled tap water for 4 minutes, strained and left to cool to 27°C. Honey (480 g) was stirred into the brewed tea. Into each preparation was mixed one sunflower (15×15 cm including petals, with a central disk of 5 cm), five bees (0.1 g each, with 0.5 cm length), and one agar plate with skin strains (between 17-20 colonies). The beverages were fermented in glass jars (Bormioli Rocco, Italy, h = 30 cm, d = 10 cm), in triplicate, each covered with a synthetic cloth. Fermentation occurred at room temperature (20±2°C). The pH and Brix of the samples were measured using a pH meter (Metrohm, model 913, Switzerland) and a refractometer (Atago Co., Ltd, Tokyo, Japan), respectively.

### Strain characterisation from hand skin

To characterise the cultivable bacterial species in the hands, different visual morphotypes were selected based on colony morphology (color, shape, elevation, pigmentation, and opacity). A representative of each morphotype was then restreaked onto a new plate, and the selected clones were mixed in a 2-ml screw-top tube containing 300 μl biomol water, 100 mg of 0.1 mm-diameter zirconium beads and 100 mg of 0.5 mm-diameter zirconium beads (Sigma, St. Louis, MO, USA). The tube was then vigorously shaken in a Precellys-24 homogenizer (Bertin Technologies, France) for 20 s at 4.5 m/s. The supernatant of this lysis was used for DNA amplification targeting the 16S rRNA gene, using the 27-F and 1492-R primers ^76^. Thermal cycling conditions applied were: (i) initial denaturation at 94 °C for 5 min; (ii) 35 amplification cycles (94°C for 40 s, 50°C for 40 s, and 68°C for 1.5 min); and (iii) final extension step at 68°C for 10 min. The PCR products were purified using the ExoSAP-IT (Thermo Fisher Scientific, Waltham, MA, USA) and the sequences were analyzed with the NCBI BLAST tool ^77^.

### Sampling of ingredients and inocula, and spontaneous beverages during fermentation

As well as characterising the species present on the fermenter’s hands, all ingredients were sampled for subsequent DNA extraction. Samples of the spontaneously fermented beverages were also collected over the course of the experiment, whenever a physical change in the beverages was observed. The first sample collection occurred after 30 days, where we observed evidence of fermentation in all three jars, from the decrease in Brix and pH. In the subsequent period, the cloth was removed, and the jars were exposed to the environment, to mimic the original experiment. We observed that some fruit flies were naturally attracted to the liquid (this is one of the limitations of our research, as this stage was not measured). After 15 days of open fermentation, we observed a biofilm starting to form on two jars, which were sampled. After 12 subsequent fermentation days (57 days in total), all batches had formed biofilms, and each batch was sampled from both the liquid and the biofilm. At this point, the biofilms were each transferred to a new jar with fresh sweetened tea prepared as above. After 9 further days of fermentation covered with the cloth (a total of 66 days), we observed the formation of cleaner biofilm layers on the top of the mothers and new samples were collected from both the biofilm and the liquid. The three spontaneous beverages were then maintained, and renewed every two weeks for 4 months, to test their stability (i.e. would they persist). Once satisfactorily demonstrated, the biofilms were combined in one jar, fermented in a new sweetened tea for 9 days, and new samples were taken from the liquid and the unified biofilm (totalling 186 days, around 6 months, from the beginning of the experiment).

Samples were named according to the following formula: S[st]_[br][s], where [st] is the sampling time (numbered 1-5, corresponding to 30, 45, 57, 66 and 186 days, respectively), [br] is the biological replicate (numbered 1-3), and [s] is the sampling state (L for liquid and M for mother).

### DNA extraction and sequencing strategies

The DNA extraction and sequencing of the initial experiment samples were performed as mentioned in Kothe et al. ^78^. The DNA of the sunflower, honeybees and green tea, as well of the samples collected during the spontaneous fermentation replication experiment (liquid and biofilm) were extracted using the Qiagen DNeasy PowerSoil Kit. For the ingredients and the biofilms, 5 g of each were mixed with 45 ml of saline water (0.9% NaCl), placed in stomacher bags (BagPage, Interscience, France) and homogenized in a laboratory blender Stomacher 400 (Seward, Struers, Denmark) at high speed, for 2 min. This mixture was filtered (filter porosity of 280 microns) and subsequently the liquid was centrifuged to concentrate the cells. For the liquids, 10 ml were centrifuged and the pellets were used for the DNA extraction.

DNA samples were sent to BGI Group (Hong Kong, China) for metagenomic sequencing using the DNBseq PE150 platform. Metagenomic reads corresponding to the *Helianthus annuus* (cultivar XRQ/B, RefSeq GCF_002127325.2), *Apis mellifera* (strain DH4, RefSeq GCF_003254395.2) and *Camellia sinensis* (cultivar Shuchazao, RefSeq GCF_004153795.1) were filtered out from the samples of sunflower, bee, and green tea, respectively, using Bowtie2 ^79^. The proportion of plant/bee reads in each sample was then assessed using Sam tools flagstat ^80^.

Since the number of reads remaining after filtering out plant and bee sequences was very low, we decided to complement our analysis with amplicon sequencing. The same DNA samples were sent to BGI Group for sequencing. Paired-end 300 sequencing was performed on the DNBSEQ platform, targeting a minimum of 30,000 reads per library. The sequencing focused on the 16S rRNA V3-V4 region using primers 338F and 806R ^81–83^, as well as the ITS2 region using primers ITS3 and ITS4 ^84,85^.

### Metagenomic analyses

Quality control and preprocessing of fastq files were performed with fastp v.0.23.2, using --cut_front --cut_tail --n_base_limit 0 --length_required 50 parameters ^86^. Paired-end reads were assembled into contigs using SPAdes v.3.15.5 with -meta and default options ^87^. We then predicted genes using Prodigal v.2.6.3 using default settings and meta procedure, and marker genes were extracted using fetchMG v.1.0 ^88,89^. Thereafter, taxonomic assignments were made using the marker gene *ychF,* whose closest homologue was assigned by a BLAST search on all available sequences in the NCBI protein database. Species composition plots were created in R v.3.6.1 using the package ggplot2 v.3.3.2.

For the ingredients, as we had low reads, we also estimated microbial composition by mapping the reads against the catalogue NCBI BLAST nr+euk in kaiju v. 1.9.0 ^90^. Amplicon data were processed with the FROGs pipeline ^91^, following the approach described by Jaeger et al. ^92^, with a filtering threshold applied to Operational Taxonomic Unit (OTU) abundances set at 0.1%.

### Safety investigation

Virulence factors was searched for in the metagenomic reads using MyDbFinder tool (https://cge.food.dtu.dk/services/MyDbFinder/) in combination with the Virulence Factor Database (VFDB) ^93^, applying 98% identity and 100% minimum length filters.

### MAGs and pangenomic analysis

Genome binning and refinement of all metagenomic sequences were performed using metawrap-mg v.1.3.2. (-maxbin2-concoct-metabat2 options). The resulting bins were refined with the bin_refinement module (-c 80 -x 5 options). For eukaryotes, we considered as MAGs of quality those bins that aligned with >60% of their sequence length to fungal reference genomes from GenBank, and completeness >80% with BUSCO v.5.5.0, dataset saccharomycetes_odb10.

To assess the genomic relatedness between *Fructobacillus* MAGs from our spontaneous beverages, all publicly available genomes of this genus were downloaded from NCBI (https://www.ncbi.nlm.nih.gov/datasets/genome/?taxon=559173). The genomes were then annotated with Prokka v.1.14.5, and the proteomes were used for pangenomic analysis. The core pangenomes for 57 *Fructobacillus* proteomes were calculated using the Bacterial Pan Genome Analysis tool (BPGA) ^94^, allowing the identification of conserved orthologs. These conserved orthologs were extracted from each genome, and their amino acid sequences were aligned and concatenated to generate a super matrix for phylogenetic analysis. A phylogenetic tree was constructed using IQtree 2 ^95^, following an automated workflow provided by the script available at https://github.com/WeMakeMolecules/Core-to-Tree, with the core_seq.txt and DATASET.xls files obtained as outputs from the BPGA run. Unique proteins identified in each cluster during pangenomic analysis were annotated using BLASTp on the NCBI server (https://blast.ncbi.nlm.nih.gov/). Average Nucleotide Identity (ANI) analysis was performed between the MAGs and the closest strains using JSpecies ^96^ to measure genetic relatedness.

### Metagenomic reads from other studies

Besides our spontaneous kombuchas, for comparison we also collected from the literature studies using shotgun metagenomics on different fermented beverages (**Table S8**). We found 51 samples of kombucha ^3,8,17,28^, 20 of wine ^29^, 13 of vinegar ^28,30^, 11 of water kefir ^28,31,32^, 7 of lambic beer ^33^, 6 of pu-erh tea ^34^, 5 of pulque ^35^, and 6 of other fermented beverages using fruits, cereals and honey ^28^.

### Principal Component Analysis (PCA)

Metagenomic reads from the literature and from this study were processed with SIMKA v.1.5.1 to generate dissimilarity matrices based on the presence and absence of shared k-mers across samples ^97^. A k-mer size of 21 was established, and a presence-absence matrix, using Bray-Curtis dissimilarity, was used to assess microbial compositional differences between samples. A PCA biplot was created using the ggplot2 package in R v.4.4.1, complemented by the ggrepel package.

#### Data availability

The raw sequences of metagenomic reads were deposited on the European Nucleotide Archive (ENA) under the BioProject ID PRJNA1192292 and the MAGs are available at https://data.mendeley.com/datasets/g29zhr8w7y/1.

### Study limitations

Our study has several limitations that should be acknowledged. First, although the presence of fruit flies (*Drosophila* spp.) was observed during the fermentation process, their potential influence on microbial dynamics was not quantitatively measured or controlled. However, it is known that fruit flies have historically been valuable in vinegar fermentation, as key sources of *Acetobacter* spp., and vectors that inoculate new batches and exchange microbial populations between fermenting batches ^72,73^. Based on these robust historical precedents, we suspect that, rather than confounding the experiment, the appearance of the fruit flies was crucial to it. Additionally, there is a notable time gap between Samplings 4 (S4) and 5 (S5), with four months elapsing without sample collection. Samples were not taken as they were not necessary to address the research question; however, this gap may have led us to miss potentially significant microbial transitions occurring during this period. Further samples could be taken to trace the intermediate changes observed between S4 and S5. Hand swab samples were analysed using only culture-dependent methods, which may have limited the detection and characterisation of the full microbial diversity present. This choice was due to the typically low microbial biomass in skin samples, which challenges DNA extraction and often leads to human DNA contamination. While more recent studies have developed methods to improve DNA extraction for skin microbiome samples ^74,75^, many still result in low DNA yields, making culture-independent approaches difficult to use. With the other samples, we did not eliminate plant or animal DNA before sequencing, which reduced the number of usable microbial reads for the ingredient samples, impacting the depth of our microbial analysis. Lastly, the pangenomic analyses were conducted only using metagenome-assembled genomes (MAGs), rather than full genomes from isolated microorganisms. Although we only used MAGS with high completeness and low contamination scores, they may not fully capture all the genes present.

## Acknowledgments

KW was employed at former restaurant Amass in Copenhagen, Denmark, where he conceived and carried out the initial spontaneous kombucha experiments. JE was funded by the Mortimer May DPhil Scholarship in Human Geography at Hertford College, Oxford, UK, during his study of KW’s initial experiments and samples. Sequencing of the initial samples was supported by M Thomas P Gilbert’s grant from the Danish National Research Foundation, grant number DNRF143. CIK, KW and JDE were funded by the Novo Nordisk Foundation, grant number NNF20CC0035580, during the replication experiment and subsequent analysis. We would also like to thank Henry Stevens, former Beverage Manager at former restaurant Amass in Copenhagen, Denmark, who made the kvass that became the bread bucha, and SFI team members for comments on the manuscript.

## Supplementary files

**Fig. S1.** Photos from the experiments. **A.** Biofilm formation around sunflowers in original experiment, 2017. **B & C.** Assembly of sunflowers, bees, and skin microbial cultures in a sweetened green tea for replication experiment, 2022.

**Fig. S2.** Principal Component Analysis (PCA) 3D plots. **A.** 3D PCA plot showing the spatial distribution of our fermented beverage samples (black points) in relation to other fermented beverages from the literature, highlighting PC3. **B.** A more focused 3D view of the PCA plot, showing the clustering differences at various fermentation stages of our samples. Interactive 3D plots are available in the supplementary materials as HTML files.

**Table S1.** Metadata of the samples used in this study (initial experiment samples, ingredients, and replication experiment samples), with information on the sampling year, fermentation time, quality control and accession numbers.

**Table S2.** Relative abundance of microbial species based on the *ychF* marker gene, calculated using sequence coverage of assembled reads. The colour code indicates abundance of each species, ranging from red (low abundance) to green (high abundance).

**Table S3.** Virulent factors identified in our spontaneous fermented beverage metagenomes.

**Table S4**. Results from culture-dependent and culture-independent analyses of the ingredients used in the replication experiment.

**Table S5.** Prokaryotic and eukaryotic MAGs recovered in this study, including quality metrics, GC content, genome size and the ANI comparisons with the closest reference species. Names highlighted in red indicate potential new species. The colour code indicates the MAGs’ completeness, ranging from low (red) to high (green), and contamination, ranging from low (red) to high (blue).

**Table S6.** Metrics of the *Fructobacillus* spp. used for the pangenomic analysis (sheet ‘Metrics’), and ANI analyses for the *F. evanidus* strains (sheet ‘ANI’). The colour code indicates the ANI values ranging from low (red) to high (green).

**Table S7.** Unique genes annotated with NCBI BLAST from bee-associated *F. evanidus* genomes available in the public database (sheet ‘bees’), and from MAGs of *F. evanidus* associated with the fermented beverages recovered in this study (sheet ‘beverage’).

**Table S8.** Details of the fermented beverage samples from the literature used to build *Fig. 5*.

