## Supplementary figures and images for "Possible origins of kombucha in spontaneous fermentation"

### Fig. S1

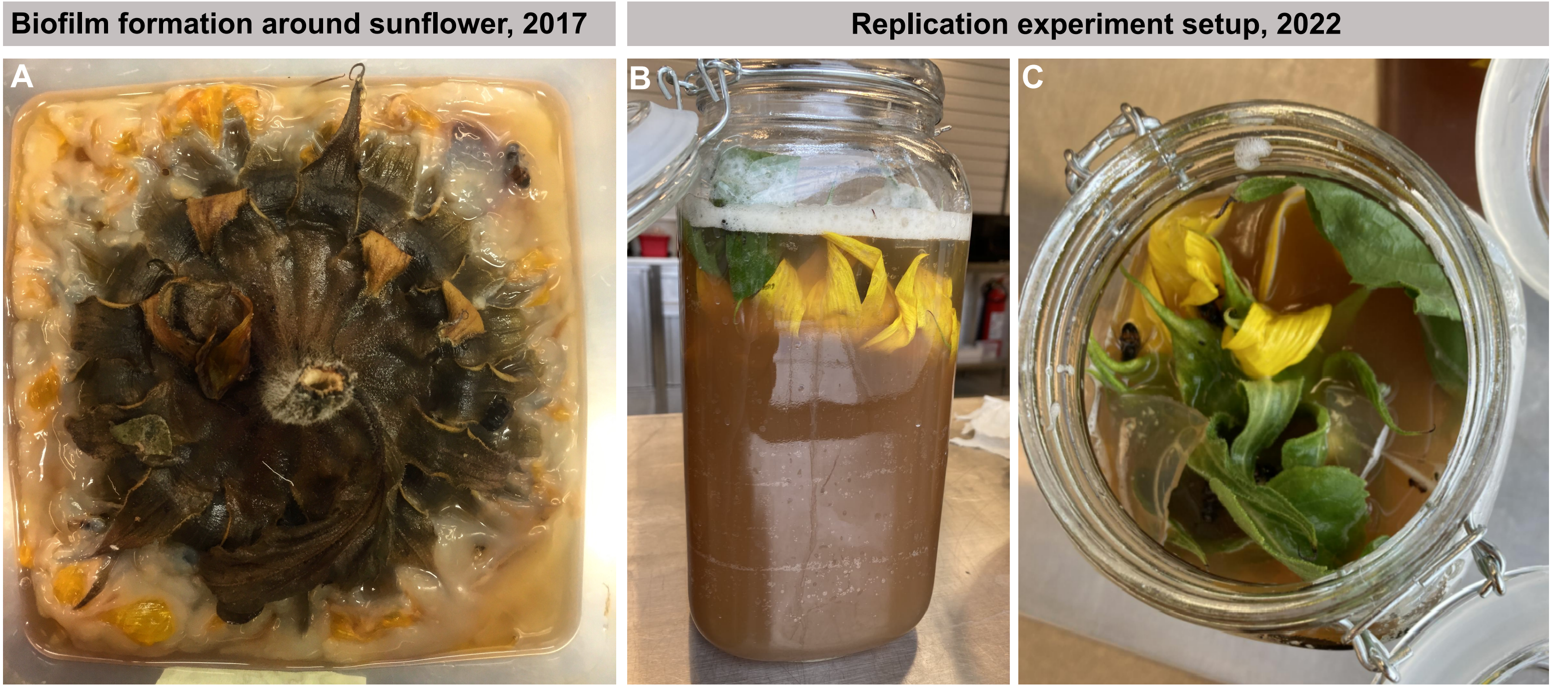

### Fig. S2

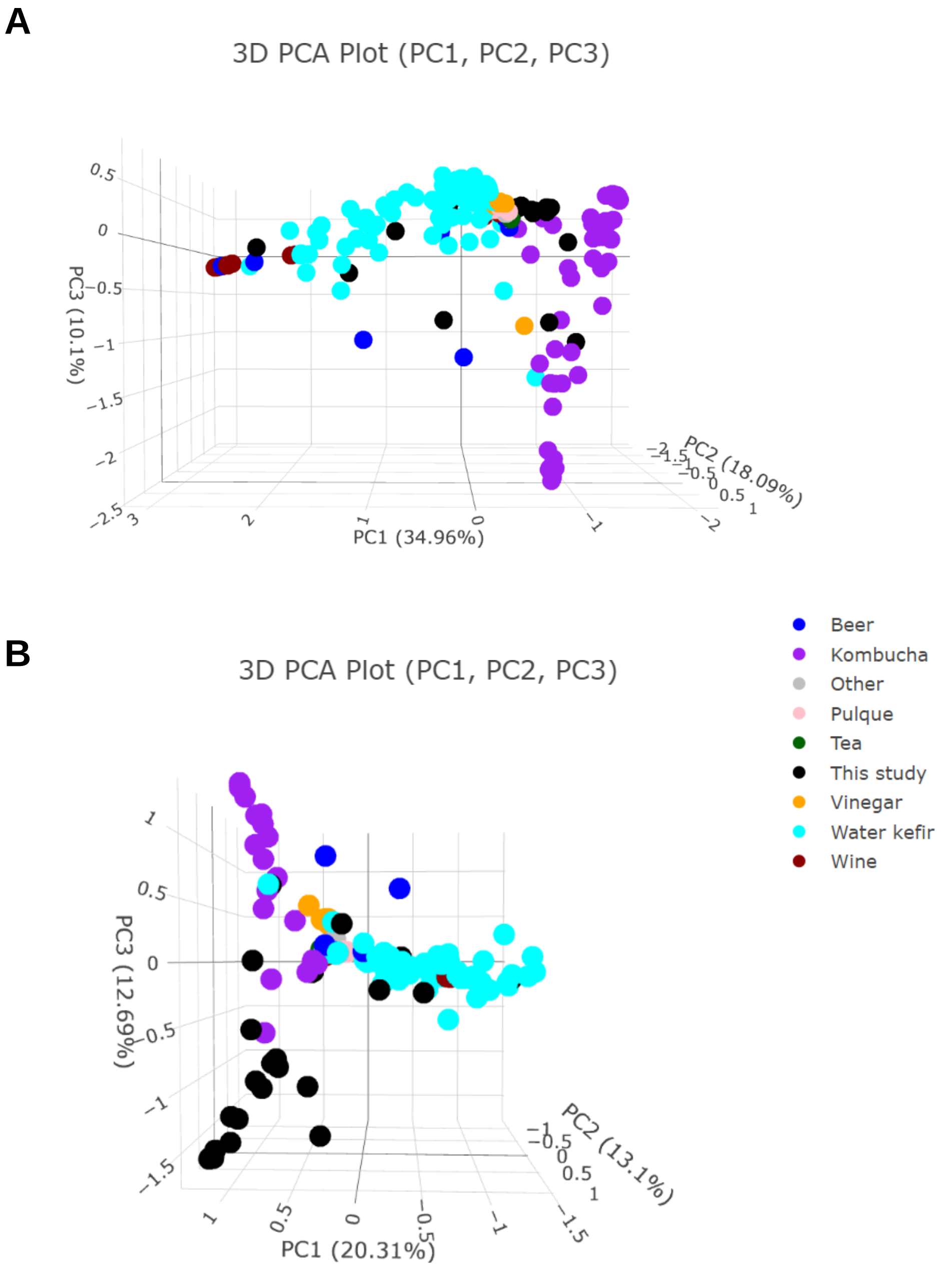
